# Crosstalkr: An open-source R package to facilitate drug target identification

**DOI:** 10.1101/2023.03.07.531526

**Authors:** Davis T. Weaver, Jacob G. Scott

**Affiliations:** Case Western Reserve University School of Medicine, Cleveland, OH, 44106, USA; Translational Hematology Oncology Research, Cleveland Clinic, Cleveland OH, 44106, USA; Department of Physics, Case Western Reserve University, Cleveland, OH, 44106, USA

## Abstract

In the last few decades, interest in graph-based analysis of biological networks has grown substantially. Protein-protein interaction networks are one of the most common biological networks, and represent the molecular relationships between every known protein and every other known protein. Integration of these interactomic data into bioinformatic pipelines may increase the translational potential of discoveries made through analysis of multi-omic datasets. Crosstalkr provides a unified toolkit for drug target and disease subnetwork identification, two of the most common uses of protein protein interaction networks. First, crosstalkr enables users to download and leverage high-quality protein-protein interaction networks from online repositories. Users can then filter these large networks into manageable subnetworks using a variety of methods. For example, network filtration can be done using random walks with restarts, starting at the user-provided seed proteins. Affinity scores from a given random walk with restarts are compared to a bootstrapped null distribution to assess statistical significance. Random walks are implemented using sparse matrix multiplication to facilitate fast execution. Next, users can perform in-silico repression experiments to assess the relative importance of nodes in their network. At this step, users can supply protein or gene expression data to make node rankings more meaningful. The default behavior evaluates the human interactome. However, users can evaluate more than 1000 non-human protein-protein interaction networks as a result of integration with StringDB. It is a free, open-source R package designed to allow users to integrate functional analysis using the protein-protein interaction network into existing bioinformatic pipelines. A beta version of crosstalkr available on CRAN (https://cran.rstudio.com/web/packages/crosstalkr/index.html).

## 1 Introduction

In the last few decades, interest in graph-based analysis of biological networks has grown substantially. Researchers have leveraged gene regulatory networks, protein-protein interaction networks, and metabolic networks to make predictions about disease gene prioritization, drug target identification, drug repurposing, and patient outcomes^1–6^. The explosion of methods and data repositories has led some to call this the “Biological Network Era”^7^. For the purpose of this paper, we will focus on genome-scale protein association networks, often called protein-protein interaction networks (PPI networks). Genome scale PPI databases attempt to represent the relationships between every known protein and every other known protein. PPI networks can be inferred through expert curation of the literature to identify empirical interactions between proteins^8^. PPI networks can also be inferred using gene or protein co-expression, gene coevolution, and other methods to reduce the false negative rate associated with expert curated PPI networks^9,10^.

Researchers have found many uses for PPI networks. For example, they have applied graph search and graph clustering algorithms to biological networks in an effort to derive disease-specific subgraphs^11–13^ or identify potential drug targets^4,6,14,15^. Further, researchers have applied simple node ranking methods to prioritize genes during the generation of predictive biomarkers^16^. Many of the key bioinformatic steps are shared between these use cases. Researchers must interface with a publicly available genome-scale interaction database, reduce the size of the total network to a manageable subnetwork, and finally score nodes so that they can be ranked.

Graph filtration (some times called graph pruning or graph reduction) is a critical step in interactomic pipelines due to the continued expansion of known protein-protein interactions^8,17^. The best available PPIs have catalogued more than 1 million individual interactions^9^. Network pruning is a key step due to the lack of context specificity in PPI repositories. All recorded interactions between proteins are made available to users, regardless of the cell cycle or phenotypic context in which they were measured. Therefore, it is critical to reduce the full PPI to a more phenotype-specific subgraph. Finally, manipulation of these data structures is extremely cumbersome and makes analysis slow. For example, the full PPI downloadable from StringDB contains 19,271 nodes and 5,934,147 edges. It requires 191.4 MB of memory.

PPI filtration is typically performed using a walk-based algorithm or through integration with additional phenotypic data such as gene expression^4,11^. One of the most well-studied algorithms in this context is random walk with restarts. Random walk with restarts (RWR) has been used and adapted across disciplines and industries for applications ranging from internet search engines to drug target identification^18–20^. There are a growing suite of tools available in R for analyzing graph-structured data^21,22^, including a few R packages that implement RWR^23,24^. These tools require some understanding of graph data structures and ask the user to find, download, and manipulate the relevant biological networks into adjacency matrices or igraph objects. In this paper, we will describe crosstalkr, a free, open-source R software package. Crosstalkr provides a streamlined interface to facilitate all three of the most common steps of an interactomic analysis. Users can interface with online PPI repositories, prune or filter the resulting networks, and rank nodes based on a variety of graph-based scoring methods. In addition, crosstalkr is optimized to facilitate one-line implementation of an RWR-based algorithm designed to identify functional subgraphs of protein-protein interaction networks (PPI)^11^. In addition to providing a clean interface for non-graph theorists, crosstalkr improves upon existing tools by implementing methods to facilitate *in silico repression*.

## 2 Design and Data Sources

Crosstalkr facilitates all 3 of the most common steps in an interactomic pipeline; interfacing with online PPI repositories, pruning the resulting network, and ranking nodes to produce scored gene sets (**Figure 1**). We implemented this functionality in the “load_ppi”, “gfilter”, and “node_repression” methods (described below).

**Figure 1.**
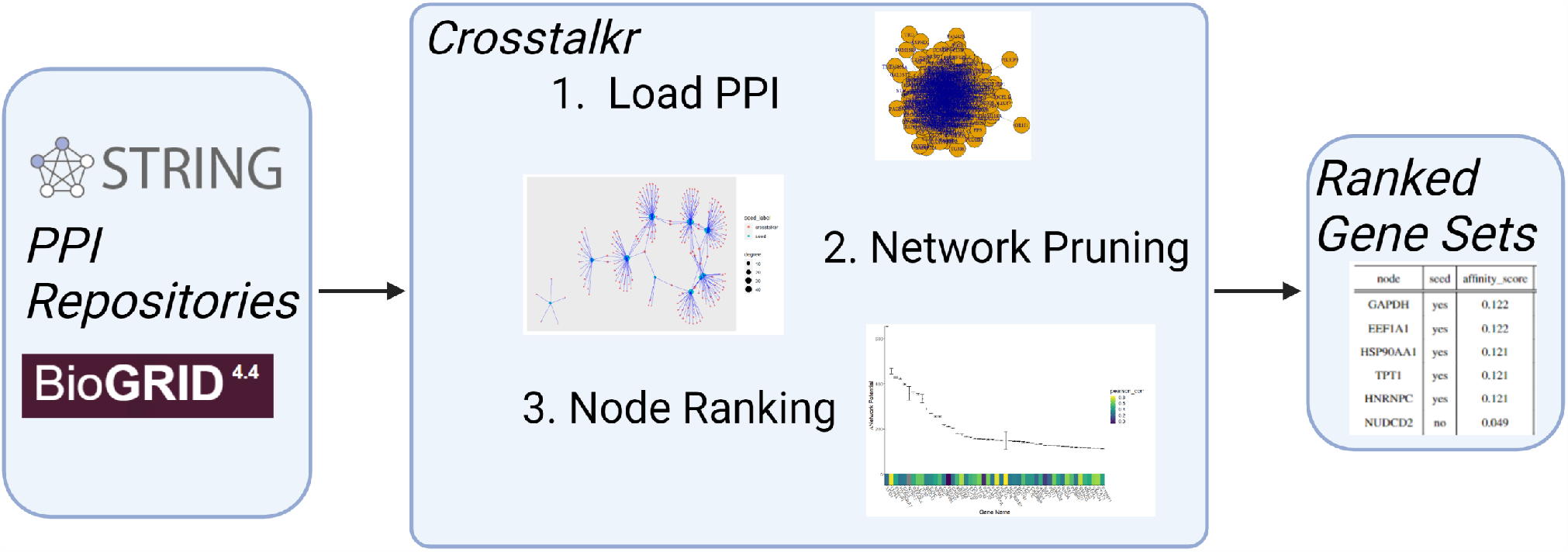
Example workflow for crosstalkr software package. crosstalkr supports all 3 of the most common steps in an interactomic pipeline; interfacing with online PPI repositories, pruning the resulting network, and ranking nodes to produce scored gene sets.

### 2.1 load_ppi

‘load_ppi’ allows users to interface directly with online protein interaction repositories. Currently, we support StringDB^9^, and Biogrid^8^. If users specify StringDB as the ppi, they can customize the incoming PPI based on species code, interaction confidence score, and interaction type. Of note, StringDB incorporates Biogrid curated interactions in their network inference process^9^. Biogrid can therefore be thought of as a more strict subset of the PPI provided by StringDB. load_ppi will standardize the incoming data and return an igraph object where proteins are vertices and binary interactions are undirected edges. However, users do not necessarily need to interact with load_ppi directly. The methods below will call load_ppi if the ‘use_ppi’ bool is set to TRUE. Finally, users are encouraged to provide a file path to the ‘cache’ argument. The returned igraph object will then be stored at the local file path as a .Rda file rather than requiring users to repeatedly download resources from online repositories. By default, cached resources will be used when load_ppi is called subsequently.

### 2.2 gfilter

‘gfilter’ and related methods allow users to reduce large graphs to subgraphs based on a user-supplied method. Users can supply a graph as an igraph object or use a PPI network. All node scoring methods in the igraph package^21^ are supported, in addition to simple node ranking based on a user-provided named numeric vector (i.e. gene expression). Node scoring methods also include one thermodynamic measure, network potential (also called Gibbs free energy)^25^. Finally, users can use an RWR-based method (described below). In all cases, users specify the number of nodes to be kept (*n*), and whether nodes should be ranked in descending order. After computing the node score based on the user-supplied method, a subgraph is creating using the ‘induced_subgraph’ igraph method.

### 2.3 compute_crosstalk

In the compute_crosstalk function, we implement a system for identification of phenotype-specific subnetworks. If users plan to search a supported protein protein interaction network, they are only required to provide a vector of seeds proteins. Seed proteins are used as the starting point for construction of the phenotype-specific sub-network, analogous to search terms in a web search. In this situation, compute_crosstalk will:

1. Download the requested PPI (or load it from the provided cache)
2. Process the requested PPI into a sparse adjacency matrix.
3. Perform a random walk with restart using the user provided seeds to generate affinity scores for every protein in the PPI.
4. Perform many random walks with restarts from n random seeds with a matching degree distribution to generate a null distribution of affinity score.
5. Compare the affinity scores to the null distribution to compute an adjusted p-value (using the method specified in p_adjust)
6. Remove proteins with an adjusted p-value < significance_level

Users can make use of caching to store processed PPIs and speed up future analyses substantially. Users can also make use of parallel computing by setting the ncores parameter > 1. For a formal definition of the algorithm, refer to **Section 5**.

### 2.4 node_repression

The node repression function provides support for *in silico* repression, a method to rank nodes for the identification of potential drug targets. *In silico* repression attempts to score the importance of a given node by computing some global measure of network state before and after node removal. A more formal definition of our *in silico* repression implementation can be found in **Section 5**. Currently, network potential (sometimes called Gibbs Free Energy) is the only state function available. Generalizing node_repression to any state function that scores nodes is an active area of development.

### 2.5 Data Sources

Users can leverage two high quality PPI networks through crosstalkr; StringDB and Biogrid^8,26^. Biogrid is the most comprehensive available database of expert-curated empirically measured protein-protein interactions. StringDB expands the coverage of Biogrid by estimating interactions using gene expression, genomic context, and other evidence streams. While Biogrid supports dozens of species, we only provide direct integration for homo sapiens. Crosstalkr provides direct access to all core species support by String.

## 3 Example Uses

Here, We present two possible bioinformatic pipelines for drug target identification. Both rely on the integration of RNA or protein expression data from a given model system with interactomic data from protein-protein interaction networks. Integration of specific phenotypic data helps address the shortcoming that publicly available PPI repositories are not context-specific^26^. In the first, we execute a two-step pipeline, where the PPI network is filtered based on RNA expression data. We then rank the nodes again based on betweenness centrality to identify the most critical hub genes of the network. In the second, we use *in silico* repression to rank the nodes rather than a simple node scoring method after a filtration step.

For these examples, we will rely on previously published gene expression data from a Ewing Sarcoma cell line (A673)g (described in more detail here). Ewing Sarcoma is a rare bone malignancy primarily seen in pediatric and adolescent patients. It is thought to be driven by a fusion event that creates an abberant EWS-FLI1 transcription factor^27^. EWS-FLI1 then activates downstream effectors that drive cellular growth and proliferation. The specific transcriptional effects have been studied extensively in the last 30 years^28–30^. Despite improved knowledge about the molecular mechanisms that cause Ewing Sarcoma, meaningful clinical improvements have not been made in decades^31^. While it is not our goal here to suggest specific targets for therapy, novel therapeutics with efficacy in Ewing Sarcoma are sorely needed. We will first isolate a single replicate for convenience and log2 normalize (*E* = *log*_2_(*E*)) the gene expression values *E*. Then we will format the expression values into a named numeric vector, the preferred format for interacting with crosstalkr.

### 3.1 Pipeline 1

#### 3.1.1 Graph Filtering

First, we will use the gene expression values to reduce the size of the PPI network we need to analyze (**Listing 1**). gfilter.value performs the following actions:

1. Download and modify PPI according to user-provided parameters (passed to load_ppi)
2. Take the maximum or minimum *n* genes according to the user provided named numeric vector (passed to the val parameter)
3. Call the igraph::induced_subgraph method to create a new igraph object that contains only the genes from step 2.
4. Add gene expression to the new graph object as an attribute to allow further manipulation.

**Listing 1.**
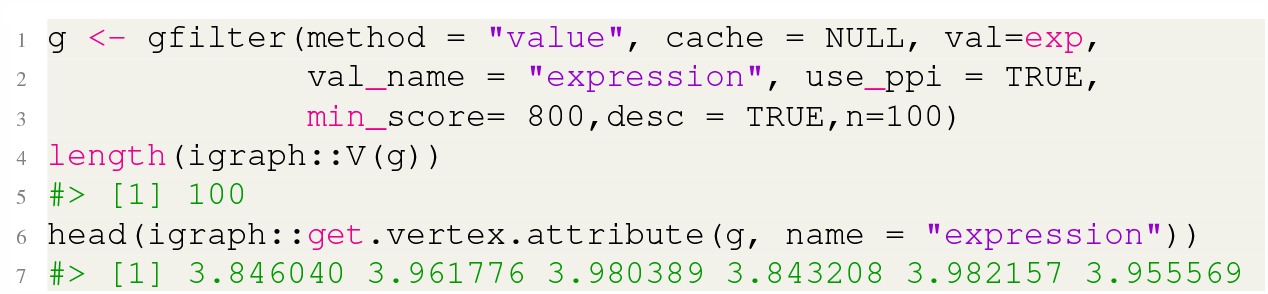
*gfilter* can reduce the size of a graph using a user-provided value to rank nodes.

#### 3.1.2 Node Scoring

Next, we will use gfilter again to score this smaller subgraph according to the betweenness centrality and return only the top 5 proteins (**Listing 2**). It is worth noting that there are a number of potential measures of centrality that could be used here, as well as many other potentially useful node scoring methods provided in the igraph package^21^.

**Listing 2.**
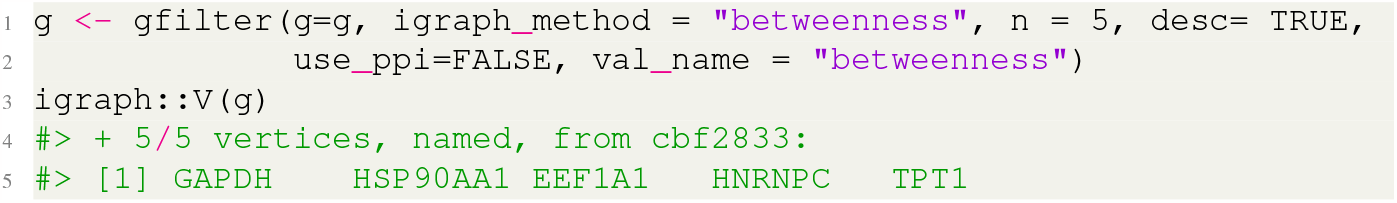
*gfilter* can return the top n nodes ranked by centrality measures like betweenness

The top 5 proteins, ranked by betweenness centrality on this reduced subgraph were GAPDH, HSP90AA1, EEF1A1, HNRNPC, and TPT1. GAPDH is a glycolytic enzyme that has also been assigned a number of other functions, and is highly expressed in normal bone marrow. HSP90AA1 encodes for a heat shock protein and has been implicated in drug resistance development in cancer. EEF1A1 is involved in protein translation. HNRNPC is a ubiquitously expressed RNA binding protein. TPT1 is a regulator of cellular proliferation, and is causally implicated in cancer development^32^. Of our 5 candidate proteins, 4 are either ubiquitously expressed housekeeping genes or proteins involved in the generic cellular stress response. Our very simple pipeline (applied to just a single sample) identified a potentially promising candidate in TPT1. TPT1 was recently associated with Ewing Sarcoma drug resistance development in the literature^28^. TPT1 was found to be significantly upregulated in a doxorubicin-resistant Ewing Sarcoma cell line compared to an embryonic fibroblast cell line.

#### 3.1.3 Disease-Specific Subnetwork Identification

Next, we use the compute_crosstalk function to identify a Ewing Sarcoma specific subnetwork from the full protein-protein interaction network (**Listing 3**). Here, we rely on the proteins identified above as seeds in the RWR-based algorithm.

**Listing 3.**
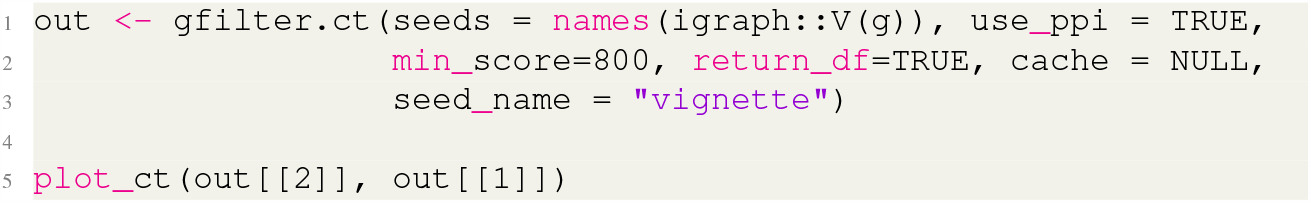
*gfilter* can reduce the size of a graph using the compute_crosstalk pipeline to rank nodes

We identified 341 proteins in the subnetwork defined by GAPDH, HSP90AA1, EEF1A1, HNRNPC, and TPT1. Most of these are neighbors of the highly connected HSP90AA1 (heat shock protein). The proteins with the highest affinity for the 5 seeds are labeled. ‘plot_ct’ is a convenience function to quickly plot the returned subgraph (**Figure 2**). Users can specify ‘prop_keep’ to improve readability by only plotting the top x% of identified proteins, ranked by affinity score. gfilter.ct also returns additional information about the proteins in the identified subnetwork (**Table 1**).

**Table 1.** top 10 proteins ranked by affinity to Ewing Sarcoma-related seeds.

| node | seed | affinity_score | adj_p_value |
| --- | --- | --- | --- |
| GAPDH | yes | 0.122 | 0 |
| EEF1A1 | yes | 0.122 | 0 |
| HSP90AA1 | yes | 0.121 | 0 |
| TPT1 | yes | 0.121 | 0 |
| HNRNPC | yes | 0.121 | 0 |
| NUDCD2 | no | 0.049 | 0 |
| TSSK6 | no | 0.049 | 0 |
| RALYL | no | 0.048 | 0 |
| POTEF | no | 0.025 | 0 |
| TOMM34 | no | 0.024 | 0 |

**Figure 2.**
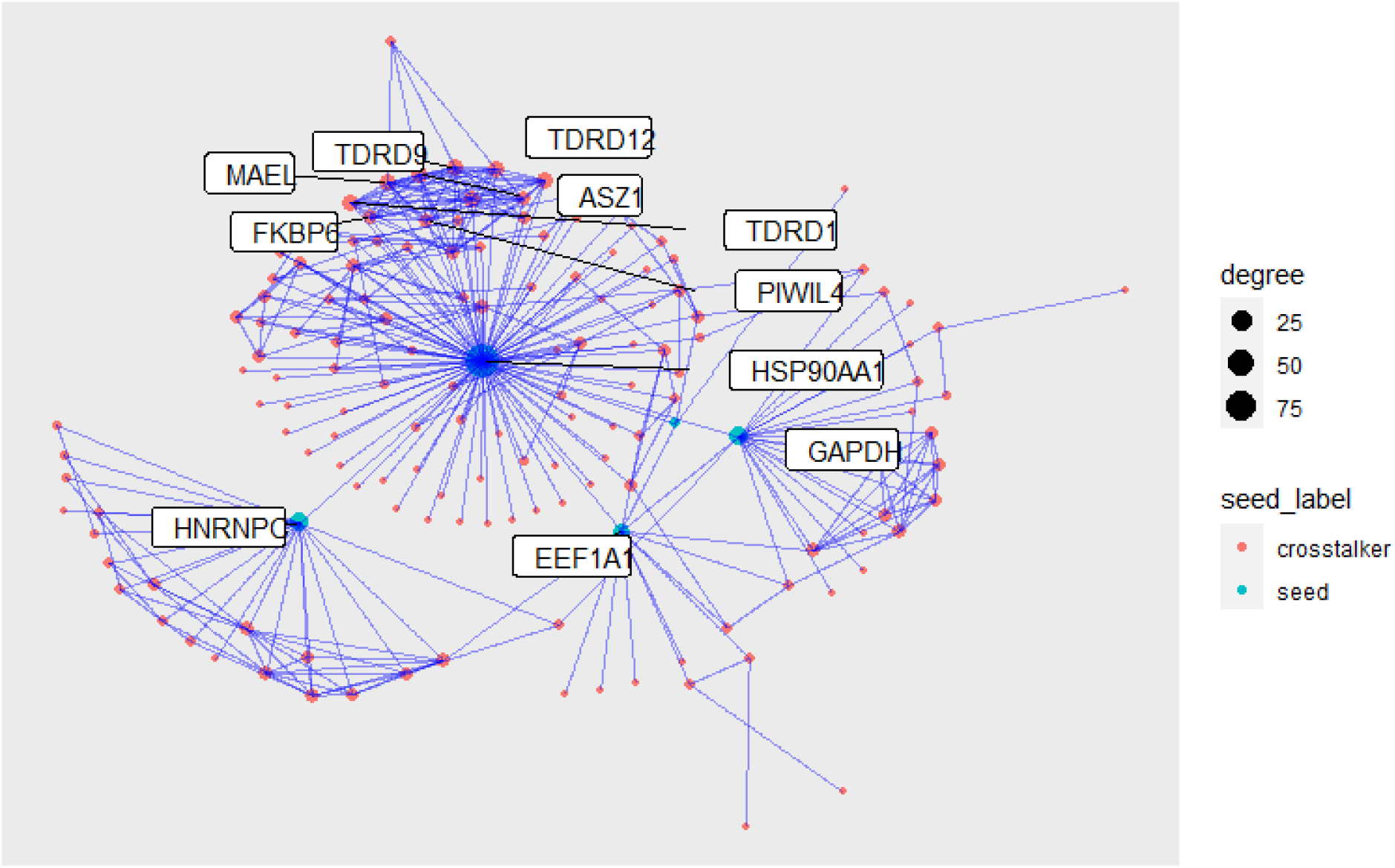
Protein-protein interaction subnetwork for GAPDH, HSP90AA1, EEF1A1,HNRNPC, and TPT1. HSP90AA1 was identified as a critical hub in the computed subnetwork.

Not including the initial seeds, NUDCD2, TSSK6m RALYL, POTEF, and TOMM34 showed the highest affinity for the provided seed proteins. Next, we will re-analyze these data using a different combination of analytic steps to illustrate more features of crosstalkr.

#### 3.2 Pipeline Two

In this pipeline, we will reduce the graph two times, first using degree (number of neighbors for a given node), and then using gene expression (**Listing 4**). The resulting graph has 500 nodes, all with both high connectivity in the PPI network and high expression in our Ewing Sarcoma cell line. There are 9624 edges (interactions) between these 500 nodes (proteins).

**Listing 4.**
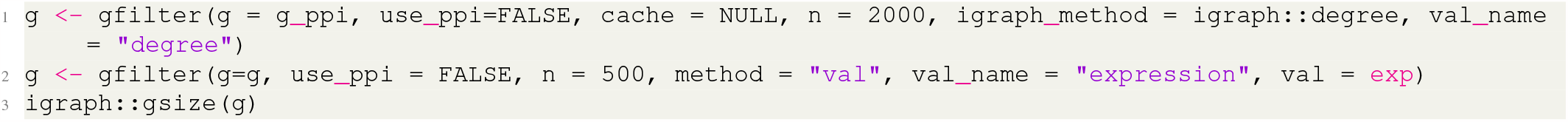
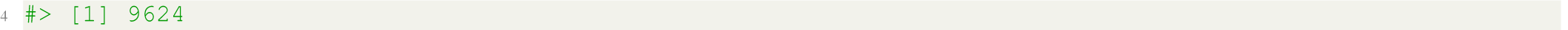
*gfilter* can reduce the size of a graph using centrality measures like degree to rank nodes.

#### 3.2.1 Node Scoring

Next, we will demonstrate a method for node scoring that uses *in silico* repression (**Listing 5**). For this example, we will use network potential as the state function. The resulting data structure is sparse matrix that shows the effect of removing the node in a given column on the state function value for the node represented in a given row. The global effect of removing a given node is then the column sum.

**Listing 5.**
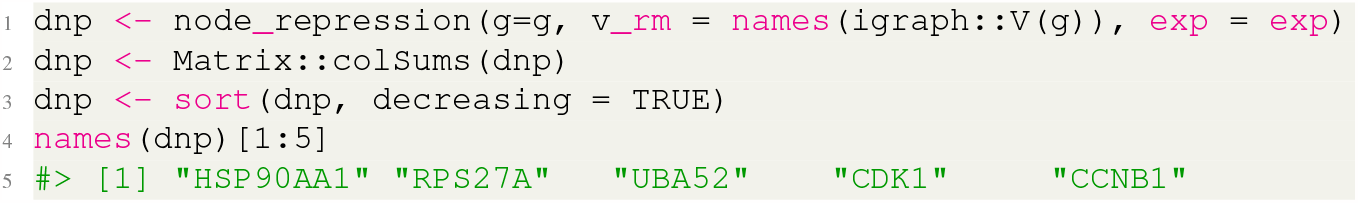
*node_repression* ranks nodes through in-silico repression.

Using this modified pipeline, we identified HSP90AA1, RPS27A, UBA52, CDK1, and CCNB1 as the most critical proteins in this Ewing Sarcoma cell signaling network. HSP90AA1 was also identified in the first instance. RPS27A is a ribosomal protein that is involved in RNA binding. UBA52 is a paralog of RPS27A and appears to be an unlikely candidate for cancer therapy. CDK1 is a cyclin dependent kinase, involved in cellular proliferation and causally associated with cancer in the literature. CCNB1 is a cyclin, a cellular regulator of mitosis.

As with the method above, our bioinformatic pipeline identified a number of ubiquitiously expressed housekeeping genes. Interactomic analyses are often biased towards proteins with extremely high connectivity (i.e. high degree). For example, HSP90AA1 was also identified as a key hub in the Ewing Sarcoma network when we analyzed these data in^6^. However, in that study, we performed a permutation test that showed the impact of HSP90AA1 was likely driven by the underlying network connectivity rather than by the actual phenotype under study. To do this, we computed a null distribution for our state function of interest by generating thousands of rewired graphs with preserved degree and then re-calculating our metric of interest. More details about this process can be found in^6^. In crosstalkr, We support this capability for the network potential example (see compute_null_dnp). Supporting this feature for additional state functions is an area of development.

## 4 Discussion

Despite the widespread integration of interactomics into bioinformatic workflows, no actively maintained R packages directly supported the core functions of an interactomic analysis. To remedy this, We developed crosstalkr, a free, open-source R package. Crosstalkr provides a toolkit for drug target identification, and supports users at every step in the pipeline. Crosstalkr ships with functions that helps users interface with centralized repositories of functional interactions, filter or prune the resulting PPI networks, and score nodes.

Many organizations have developed web applications to help industry and academic stakeholders perform basic interactomic analyses, including simple queries of protein neighborhood and random-walk inspired tests to identify functionally related proteins^11,23,26,33,34^. Unfortunately, closed-source web applications are often limited by the lack of programmatic access to the resources. We developed crosstalkr as an R package to enable rapid integration into modular, R-native bioinformatic pipelines. Further, users cannot rely on long-term maintenance of products in this sector. The vast majority of web applications (and software packages) that have been developed to assist with interactomic analyses fall into disrepair within a few years of their release^35–37^. Against that backdrop, open-source frameworks that can be extended and maintained by third-parties are far more likely to provide to provide long-term value to the academic community.

Overall, crosstalkr builds on a vibrant landscape of tools to help users interact with functional interaction networks, and provides a novel implementation of *in silico* repression. Crosstalkr has been downloaded more than 1700 times and has more than 10 stars on github, highlighting the potential impact of this work. Crosstalkr can be download directly from CRAN using “install.packages(“crosstalkr”)” from any R environment. The most recent developmental version can be found on github (https://github.com/DavisWeaver/crosstalkr).

## 5 Algorithm Definitions

### 5.1 Random Walks with Restarts

As mentioned above, compute_crosstalk is an implementation of the random walk with restarts, followed by a permutation test for signficance. The mathematical definition for a random-walk with restarts has been written many times, and can be found at^18^. Here, we provide pseudocode to describe the algorithm implementation in crosstalkr. *w* is a square adjacency matrix describing the graph. *seeds* is a vector describing the indexes of the provided seeds. *gamma* is a number describing the discount rate. *eps* describes the stop condition. When the change in p in a single iteration is less than *eps*, the algorithm will stop. *tmax* describes the largest number of iterations that the algorithm will attempt. The ouput is *p*, a vector containing the affinity score of all nodes in *w* relative to the provided *seeds*.

**Listing 6.**
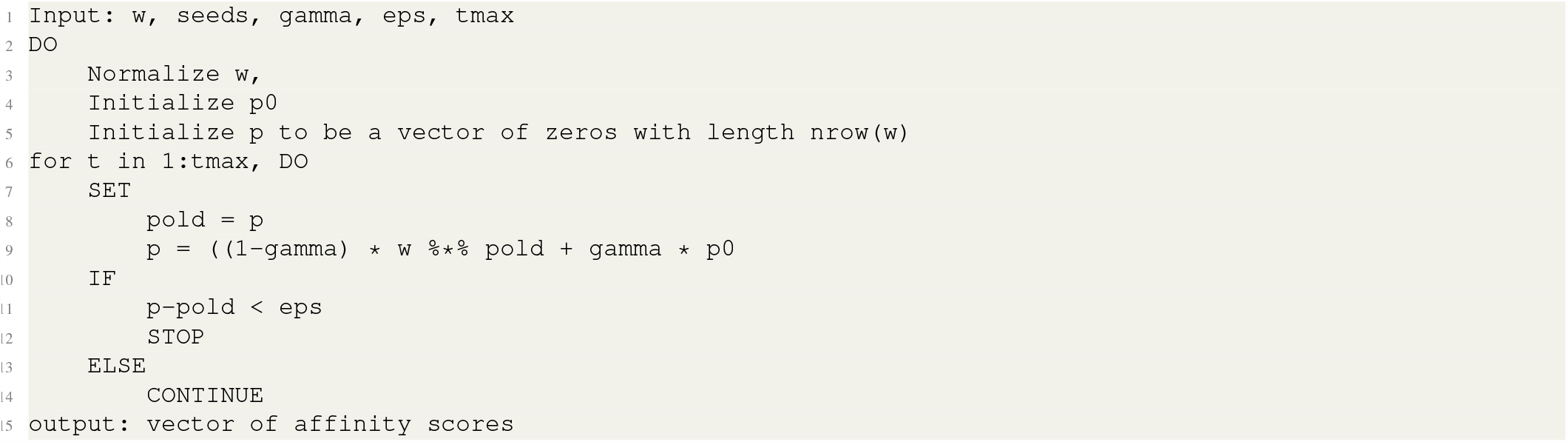
psuedocode for crosstalkr implementation of random walk with repeats

### 5.2 Bootstrapped Null Distributions

We computed bootstrapped null distributions using two related methods. In the algorithm implemented by compute_crosstalk, we bootsrapped a null by re-calculating affinity scores (through RWR) associated with a series of randomly selected seeds that shared a similar degree distribution compared to the user-provided seeds. For more details on this method, refer to^11^. In the algorithm implemented by compute_null_dnp, we bootstrapped a null by generating *n* completely new graphs with preserved degree distribution using the “keeping_degseq” method. We described this method in detail in our previous work^6^.

### 5.3 Network Potential

Network Potential (i.e. Gibbs Free Energy) is a node-specific metric that depends on the local context of each node in a graph. We discuss network potential in detail in our previous paper^6^. For a given node *V*_*i*_ and node weight (*C*_*i*_) in a graph *G*(*V, E*), network potential (*G*_*i*_) can be computed as follows:

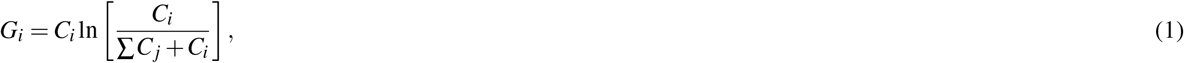

where *C*_*j*_ refers to the node weights for all neighbors of *V*_*i*_.

### 5.4 Betweenness Centrality

Betweenness centrality measures the number of shortest paths that traverse a given node *v*. Betweenness can be computed as follows:

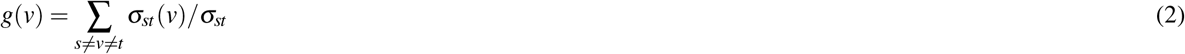

where *σ*_*st*_ is the number of shortest paths from node *s* to node*t* and *σ*_*st*_(*v*) is the number of shortest paths from node *s* to node *t* that traverse node *v*.

### 5.5 In silico repression

*In silico* repression attempts to quantify the importance of a given node *v* in some graph *G*(*V, E*) by computing the a global measure of graph state S before and after the removal of node *v* from the graph. In crosstalkr, *in silico* repression is a 6-step process:

1. Calculation of a node score *s*_*v*_ for all nodes *v ∈ V* in the graph using some state function *s*(*v*).
2. Calculation of the total network state *S* = Σ*s*_*v*_
3. Removal of a given node *i* from the graph.
4. Re-calculation of node score *s*_*i*_*v* for all nodes *v ∈ V*. (In practice, we only re-calculate the node score of those nodes affected by the removal of node *i*)
5. Re-calculation of total network state *S*_*i*_ = Σ_*i*≠*v*_*s*_*iv*_
6. Calculation of ∆*S*_*i*_ = *S −S*_*i*_ where ∆*S*_*i*_ is used to score and rank nodes.

## Availability and Requirements

- Project Name: crosstalkr
- Project home page: https://github.com/DavisWeaver/crosstalkr
- Operating Systems: Platform indpendent
- Programming language: R
- Other Requirements: R >= 2.10
- License: GPL (>= 3)

The most recent stable build can be installed directly from CRAN.

## Declarations

### Ethics Approval and consent to participate

Not Applicable

### Consent for Publication

Not Applicable

### Availability of data and materials

See Availability and Requirements section

### Author Contributions

DTW identified the area of need, wrote the software, wrote the manuscript, and edited the manuscript. JGS provided mentorship, wrote the manuscript, and edited the manuscript.

## Acknowledgements

We acknowledge contributions from Dr. Mark Chance and Dr. Mehmet Koyuturk. We would also like to acknowledge the dependencies that enable crosstalkr^21,38–48^. We would like to acknowledge Dr. Rowan Barker-Clarke and Arda Durmaz for testing the code. Finally, we would like to thank Dr. Raoul Wadhwa for providing C++ expertise.

## References

1. Shahreza, M. L., Ghadiri, N., Mousavi, S. R., Varshosaz, J. & Green, J. R. A review of network-based approaches to drug repositioning. Briefings Bioinforma. 19, 878–892, DOI: 10.1093/bib/bbx017 (2017).

2. Arrell, D. K. & Terzic, A. Network Systems Biology for Drug Discovery. Clin. Pharmacol. & Ther. 88, 120–125, DOI: 10.1038/clpt.2010.91 (2010). _eprint: https://onlinelibrary.wiley.com/doi/pdf/10.1038/clpt.2010.91.

3. Chasman, D., Fotuhi Siahpirani, A. & Roy, S. Network-based approaches for analysis of complex biological systems. Curr. Opin. Biotechnol. 39, 157–166, DOI: 10.1016/j.copbio.2016.04.007 (2016).

4. Rietman, E. A., Scott, J. G., Tuszynski, J. A. & Klement, G. L. Personalized anticancer therapy selection using molecular landscape topology and thermodynamics. Oncotarget 8, 18735–18745, DOI: 10.18632/oncotarget.12932 (2016).

5. Rietman, E. A., Scott, J. G., Tuszynski, J. A. & Klement, G. L. Personalized anticancer therapy selection using molecular landscape topology and thermodynamics. Oncotarget 8, 18735–18745, DOI: 10.18632/oncotarget.12932 (2017).

6. Weaver, D. T. et al. Network potential identifies therapeutic miRNA cocktails in Ewing sarcoma. PLOS Comput. Biol. 17, e1008755, DOI: 10.1371/journal.pcbi.1008755 (2021). Publisher: Public Library of Science.

7. Koutrouli, M., Karatzas, E., Paez-Espino, D. & Pavlopoulos, G. A. A Guide to Conquer the Biological Network Era Using Graph Theory. Front. Bioeng. Biotechnol. 8 (2020).

8. Oughtred, R. et al. The BioGRID database: A comprehensive biomedical resource of curated protein, genetic, and chemical interactions. Protein Sci. : A Publ. Protein Soc. 30, 187–200, DOI: 10.1002/pro.3978 (2021).

9. Szklarczyk, D. et al. STRING v11: Protein-protein association networks with increased coverage, supporting functional discovery in genome-wide experimental datasets. Nucleic Acids Res. 47, D607–D613, DOI: 10.1093/nar/gky1131 (2019).

10. Kim, C. Y. et al. HumanNet v3: an improved database of human gene networks for disease research. Nucleic Acids Res. 50, d632–d639, DOI: 10.1093/nar/gkab1048 (2021).

11. Nibbe, R. K., Koyutürk, M. & Chance, M. R. An Integrative -omics Approach to Identify Functional Sub-Networks in Human Colorectal Cancer. PLOS Comput. Biol. 6, e1000639, DOI: 10.1371/journal.pcbi.1000639 (2010). Publisher: Public Library of Science.

12. Chitra, U., Park, T. Y. & Raphael, B. J. NetMix2: Unifying Network Propagation and Altered Subnetworks. In Research in Computational Molecular Biology, 193–208, DOI: 10.1007/978-3-031-04749-7_12 (2022).

13. Pfeifer, B., Secic, A., Saranti, A. & Holzinger, A. GNN-SubNet: disease subnetwork detection with explainable Graph Neural Networks. bioRxiv 2022.01.12.475995, DOI: 10.1101/2022.01.12.475995 (2022).

14. Martínez, V., Navarro, C., Cano, C., Fajardo, W. & Blanco, A. DrugNet: Network-based drug–disease prioritization by integrating heterogeneous data. Artif. Intell. Medicine 63, 41–49, DOI: 10.1016/j.artmed.2014.11.003 (2015).

15. Rietman, E. A. et al. Using the Gibbs Function as a Measure of Human Brain Development Trends from Fetal Stage to Advanced Age. Int. J. Mol. Sci. 21, 1116, DOI: 10.3390/ijms21031116 (2020). Number: 3 Publisher: Multidisciplinary Digital Publishing Institute.

16. Scott, J. G. et al. A genome-based model for adjusting radiotherapy dose (GARD): a retrospective, cohort-based study. The Lancet Oncol. 18, 202–211, DOI: 10.1016/S1470-2045(16)30648-9 (2017).

17. Razick, S., Magklaras, G. & Donaldson, I. M. iRefIndex: A consolidated protein interaction database with provenance. BMC Bioinforma. 9, 405, DOI: 10.1186/1471-2105-9-405 (2008).

18. Tong, H., Faloutsos, C. & Pan, J.-y. Fast Random Walk with Restart and Its Applications. In Sixth International Conference on Data Mining (ICDM’06), 613–622, DOI: 10.1109/ICDM.2006.70 (2006). ISSN: 2374-8486.

19. Bianchini, M., Gori, M. & Scarselli, F. Inside PageRank. ACM Transactions on Internet Technol. 5, 92–128, DOI: 10.1145/1052934.1052938 (2005).

20. Navarro, C., Martínez, V., Blanco, A. & Cano, C. ProphTools: general prioritization tools for heterogeneous biological networks. GigaScience 6, 1–8, DOI: 10.1093/gigascience/gix111 (2017).

21. details, S. A. f. i. a. igraph: Network Analysis and Visualization (2022).

22. Gatto, L. & Christoforou, A. Using R and Bioconductor for proteomics data analysis. Biochimica et Biophys. Acta (BBA) - Proteins Proteomics 1844, 42–51, DOI: 10.1016/j.bbapap.2013.04.032 (2014). ArXiv:1305.6559 [q-bio].

23. Fang, H. & Gough, J. The ‘dnet’ approach promotes emerging research on cancer patient survival. Genome Medicine 6, 64, DOI: 10.1186/s13073-014-0064-8 (2014).

24. Valentini, G. RANKS: Ranking of Nodes with Kernelized Score Functions (2022).

25. Rietman, E. & Tuszynski, J. A. Using Thermodynamic Functions as an Organizing Principle in Cancer Biology. In Theoretical and Applied Aspects of Systems Biology, 139–157, DOI: 10.1007/978-3-319-74974-7_8 (Springer, Cham, 2018).

26. Szklarczyk, D. et al. The STRING database in 2021: customizable protein-protein networks, and functional characterization of user-uploaded gene/measurement sets. Nucleic Acids Res. 49, D605–D612, DOI: 10.1093/nar/gkaa1074 (2021).

27. Riggi, N. et al. EWS-FLI-1 modulates miRNA145 and SOX2 expression to initiate mesenchymal stem cell reprogramming toward Ewing sarcoma cancer stem cells. Genes & development 24, 916–32, DOI: 10.1101/gad.1899710 (2010).

28. Yakushov, S. et al. Identification of Factors Driving Doxorubicin-Resistant Ewing Tumor Cells to Survival. Cancers 14, 5498, DOI: 10.3390/cancers14225498 (2022).

29. Bennani-Baiti, I. M., Machado, I., Llombart-Bosch, A. & Kovar, H. Lysine-specific demethylase 1 (LSD1/KDM1A/AOF2/BHC110) is expressed and is an epigenetic drug target in chondrosarcoma, Ewing’s sarcoma, osteosarcoma, and rhabdomyosarcoma. Hum. pathology 43, 1300–7, DOI: 10.1016/j.humpath.2011.10.010 (2012).

30. Hameiri-Grossman, M. et al. The association between let-7, RAS and HIF-1 in Ewing Sarcoma tumor growth. Oncotarget 6, 33834–48, DOI: 10.18632/oncotarget.5616 (2015).

31. Esiashvili, N., Goodman, M. & Marcus, R. B. Changes in Incidence and Survival of Ewing Sarcoma Patients Over the Past 3 Decades. J. Pediatr. Hematol. 30, 425–430, DOI: 10.1097/MPH.0b013e31816e22f3 (2008).

32. Tate, J. G. et al. COSMIC: the Catalogue Of Somatic Mutations In Cancer. Nucleic Acids Res. 47, D941–D947, DOI: 10.1093/nar/gky1015 (2019).

33. Vella, D. et al. MTGO: PPI Network Analysis Via Topological and Functional Module Identification. Sci. Reports 8, 5499, DOI: 10.1038/s41598-018-23672-0 (2018).

34. Warde-Farley, D. et al. The GeneMANIA prediction server: biological network integration for gene prioritization and predicting gene function. Nucleic Acids Res. 38, w214–w220, DOI: 10.1093/nar/gkq537 (2010).

35. Kamburov, A., Stelzl, U., Lehrach, H. & Herwig, R. The ConsensusPathDB interaction database: 2013 update. Nucleic Acids Res. 41, d793–d800, DOI: 10.1093/nar/gks1055 (2012).

36. Kennedy, S. A. et al. Extensive rewiring of the EGFR network in colorectal cancer cells expressing transforming levels of KRASG13D. Nat. Commun. 11, 499, DOI: 10.1038/s41467-019-14224-9 (2020). Number: 1 Publisher: Nature Publishing Group.

37. Vinayagam, A. et al. Controllability analysis of the directed human protein interaction network identifies disease genes and drug targets. Proc. Natl. Acad. Sci. 113, 4976–4981, DOI: 10.1073/pnas.1603992113 (2016). Publisher: Proceedings of the National Academy of Sciences.

38. Wickham, H., François, R., Henry, L., Müller, K. & RStudio. dplyr: A Grammar of Data Manipulation (2022).

39. magrittr), S. M. B. O. a. a. c. o., Wickham, H., Henry, L. & RStudio. magrittr: A Forward-Pipe Operator for R (2022).

40. Hester, J. et al. withr: Run Code ‘With’ Temporarily Modified Global State (2022).

41. Bates, D. et al. Matrix: Sparse and Dense Matrix Classes and Methods (2022).

42. Wickham, H. et al. readr: Read Rectangular Text Data (2022).

43. Wickham, H., Girlich, M. & RStudio. tidyr: Tidy Messy Data (2022).

44. Daniel, F., Ooi, H., Calaway, R. Microsoft & Weston, S. foreach: Provides Foreach Looping Construct (2022).

45. Daniel, F., Corporation, M., Weston, S. & Tenenbaum, D. doParallel: Foreach Parallel Adaptor for the ‘parallel’ Package (2022).

46. Rainer, J., Gatto, L. & Weichenberger, C. X. ensembldb: an R package to create and use Ensembl-based annotation resources. Bioinformatics 35, 3151–3153, DOI: 10.1093/bioinformatics/btz031 (2019).

47. Xie [aut, Y. et al. knitr: A General-Purpose Package for Dynamic Report Generation in R (2022).

48. Wickham, H. et al. ggplot2: Create Elegant Data Visualisations Using the Grammar of Graphics (2022).

